# Engineering of *Pseudomonas putida* KT2440 for Broad-Chain-Length 3-Hydroxy Fatty Acid Biosynthesis

**DOI:** 10.1101/2025.11.04.686531

**Authors:** Hao Meng, Tobias Schwanemann, Marie K. Lipa, Carina V. Michel, Jialiang Xia, Lars M. Blank

## Abstract

3-Hydroxy fatty acids (3-HFAs) are versatile intermediates for bio-based polymers, fuels, and surfactants, and are advancing circular economy manufacturing. We built an acyl– CoA ligase-deficient chassis of *Pseudomonas putida* KT2440, thereby blocking 3-HFA activation and preventing its use as carbon source. This strategy decoupled synthesis from catabolism and enabled 3-HFA accumulation. We then compared two different routes for modifying the free 3-HFA composition. 1. In a direct route, overexpressing native PhaG produced C8, C10, C12, and C14 3-HFAs, achieving a total titer of 0.73 g/L in shake flasks. Functional analyses under our tested conditions support a re-assignment of PhaG: rather than acting primarily as a 3-hydroxyacyl-ACP:CoA transacylase, it functions mainly as a thioesterase, liberating free 3-HFAs from hydroxyacyl-ACP. 2. In an indirect route, overexpressing RhlA variants generated hydroxyalkanoyl-alkanoates (HAAs) that were converted to free 3-HFAs by endogenous esterase(s): RhlA from *Pseudomonas aeruginosa* PAO1 favored C8–C12 and yielded 0.47 g/L 3-HFAs, whereas RhlA from *Burkholderia plantarii* PG1 favored C10–C14 with 0.14 g/L, of which 80% was C14. Finally, we demonstrated process feasibility by up-scaling the PhaG pathway in a stirred-tank reactor. These results establish modular, stable, chassis-compatible routes for tailoring 3-HFA chain-length distributions, thereby providing a foundation for scalable, bio-based monomer supply in a circular economy.

## 1. Introduction

Given the growing concerns regarding environmental sustainability, it is imperative to actively promote the use of biodegradable polymers as a sustainable alternative to fossil-based materials. Medium-chain-length polyhydroxyalkanoates (mcl-PHA), the naturally produced polyesters stored as granules in many bacteria, have been extensively studied for their potential use as biodegradable polymers to replace petrochemical-based products (Choi *et al*., 2020; Mezzina *et al*., 2021; Muigano *et al*., 2025; Park *et al*., 2024). Unlike the intracellularly produced mcl-PHA, 3-hydroxy fatty acids (3-HFAs), a precursor of PHAs (Wang *et al*., 2012), can be easily excreted and offer more flexibility in re-polymerization or esterification (Nanthachai *et al*., 2025). Notably, 3-HFAs serve as chiral building blocks for the synthesis of pharmaceuticals, antibiotics, vitamins, fragrances, aromatics, and pheromones (Braga and Belo, 2024; Chen and Wu, 2005; Martin *et al*., 2013; Ren *et al*., 2010). Moreover, 3-HFAs can also be chemically or electrochemically converted to biofuels (Mensah *et al*., 2020; Meyers *et al*., 2019; vom Stein *et al*., 2014; Zhang *et al*., 2009), providing new alternatives to fossil fuels.

Since the chemical synthesis of chiral 3-HFAs is difficult and economically unfeasible (Jaipuri *et al*., 2004), researchers have turned their attention to the biological production. Initially, the focus was on depolymerizing the mcl-PHA (Gangoiti *et al*., 2010; Ruth *et al*., 2007; Yuan *et al*., 2008), or converting fatty acids by strains with defective β-oxidation (Chung *et al*., 2013; Chung *et al*., 2009; Park *et al*., 2015; Seto *et al*., 2010). Recently, 18 g/L of (R)-3-hydroxydecanoic acid was produced by chemically hydrolyzing the R-3-(R-3-hydroxyalkanoyloxy) alkanoic acids (HAAs), which is synthesized by acyltransferase RhlA in the *Pseudomonas aeruginosa* PAO1 strain using palm oil as a carbon source (Wang *et al*., 2025). However, that strain is recognized as a human pathogen and therefore unsuitable for industrial-scale deployment; a safe, non-pathogenic alternative chassis is required.

*Pseudomonas putida* KT2440 (*P. putida*) is frequently chosen for fatty acid-related bioproduction because it is an HV1 classified (Kampers *et al*., 2019), non-pathogenic strain with robust, versatile lipid metabolism, supported by extensive literature demonstrating its utility (de Lorenzo *et al*., 2024; Martínez-García and de Lorenzo, 2024; Nikel and de Lorenzo, 2018). Particularly, the Entner–Doudoroff (ED) pathway can generate excess NADPH, which supports NADPH-dependent biosynthesis (e.g., fatty acid synthesis) and strengthens redox defenses against fatty-acid–induced stress (Nikel *et al*., 2015; Sathesh-Prabu *et al*., 2025; Vogeleer *et al*., 2024). Under nitrogen limitation, mcl-PHA are natively synthesized from sugars, polyols, or aromatics (Beckers *et al*., 2016; Borrero-de Acuña *et al*., 2014; Salvachúa *et al*., 2020). By deleting two alcohol dehydrogenases, *pedF* and *adhP*, *P. putida* is capable of producing C6-C10 fatty alcohols (Sarwar *et al*., 2022). To interrogate the genetic basis of fatty acid metabolism, Thompson *et al*. (2020) applied the RB-Tn-Seq technology and developed an evidence-based understanding of the enzymes and pathways (Thompson *et al*., 2020). Suggested by the fitness scores, three CoA ligase genes, PP_0763, PP_4549, and PP_4550, were identified that may play a key role in activating medium-chain-length fatty acids. These genes were subsequently deleted in *P. putida,* and their functionality was demonstrated by growing the deficient strains on even-numbered chain length fatty acids (C6-C16) (Valencia *et al*., 2022). Additionally, the mutant was engineered for the production of chain length-tailored free fatty acids (titers up to 0.67 g/L) and fatty acid methyl esters.

Here, the targeted deletion of acyl-CoA ligase encoding genes that block fatty-acid activation inspired a novel strategy for 3-HFA production in *P. putida*. Deleting the respective ligases was expected to block 3-HFA activation, thereby creating a chassis that was unable to consume 3-HFAs. Accordingly, combinations of ligase deletions were tested across various chain lengths and a non-consuming chassis was obtained. This chassis then enabled broad-chain-length 3-HFA production by two different routes: (i) direct release of the carbon chain from fatty acid biosynthesis via constitutive *phaG* expression, supporting the function that PhaG was identified as a 3-hydroxyacyl-ACP:CoA thioesterase (Guss *et al*., 2025; Wang *et al*., 2012); (ii) heterologous *rhlA*-driven synthesis of HAAs followed by native hydrolysis. This study make use of still-non-resolved native HAA hydrolysis and demonstrates 3-HFA production in *P. putida* KT2440. Detailed physiological characterization and production from mL to liter-scale support our understanding of native PHA formation and β-oxidation. Overall, this study progresses toward the sustainable production of hydroxy fatty acids and related products.

## 2. Materials and Methods

### 2.1. Bacterial strains, chemicals, and DNA manipulations

The bacterial strains and plasmids used in this study are listed in Tables 1 and 2. Deletion mutants were constructed using the optimized I-*Sce*I-mediated one plasmid system (Meng *et al*., 2024) and chromosomal integrations were carried out using mini-Tn7 (Zobel *et al*., 2015). The oligonucleotides used in this study are listed in Table S1. *E. coli* PIR2 was used as a cloning host for plasmid construction, and also as a donor strain during parental mating for plasmid delivery. The *E. coli* HB101 strain was used as a helper strain for all the parental mating, whereas the *E. coli* DH5α λ*pir* pTNS1 was additionally used in the mating for Tn7 integration.

**Table 1.**
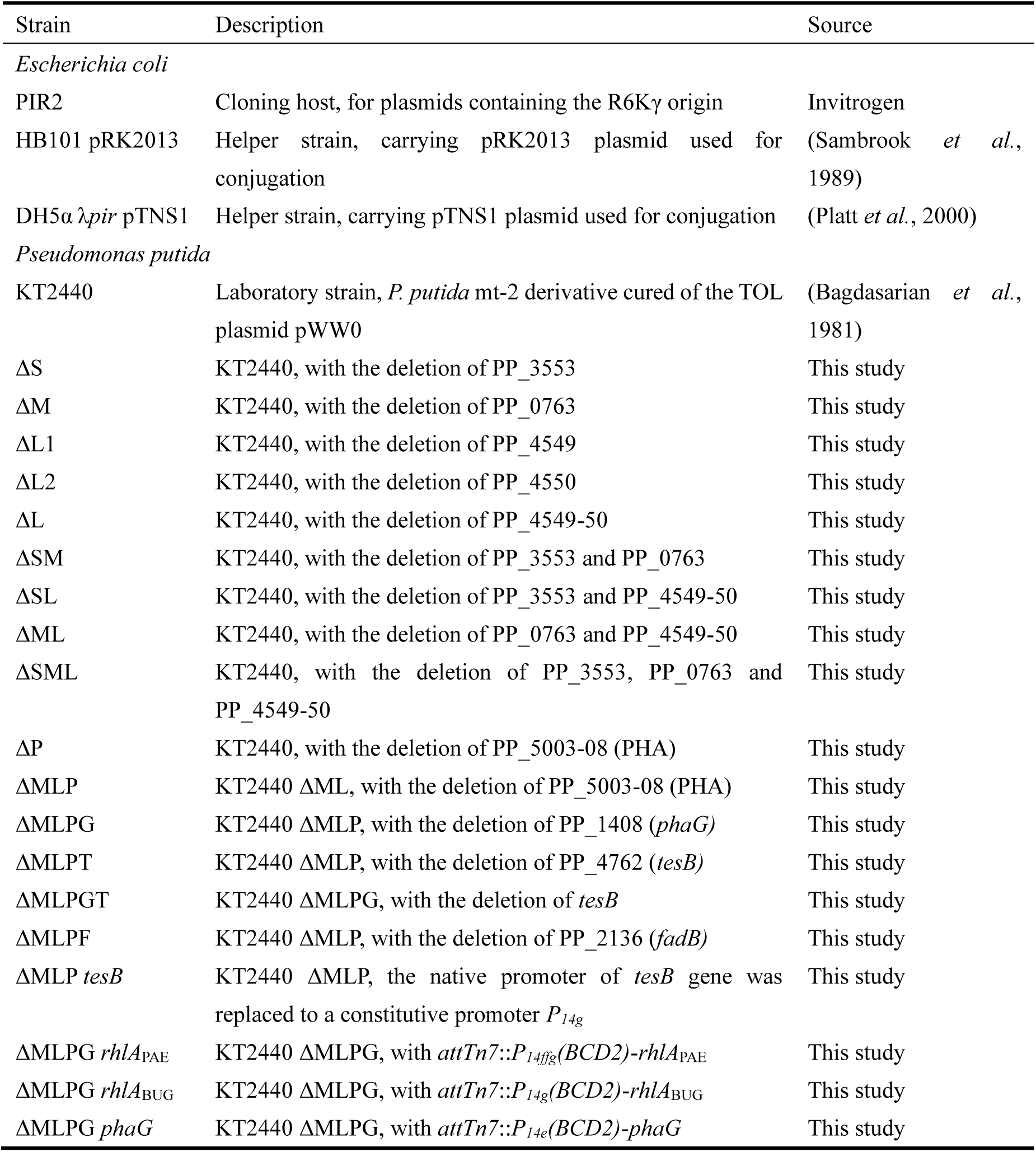
Bacterial strains used in this study.

**Table 2.**
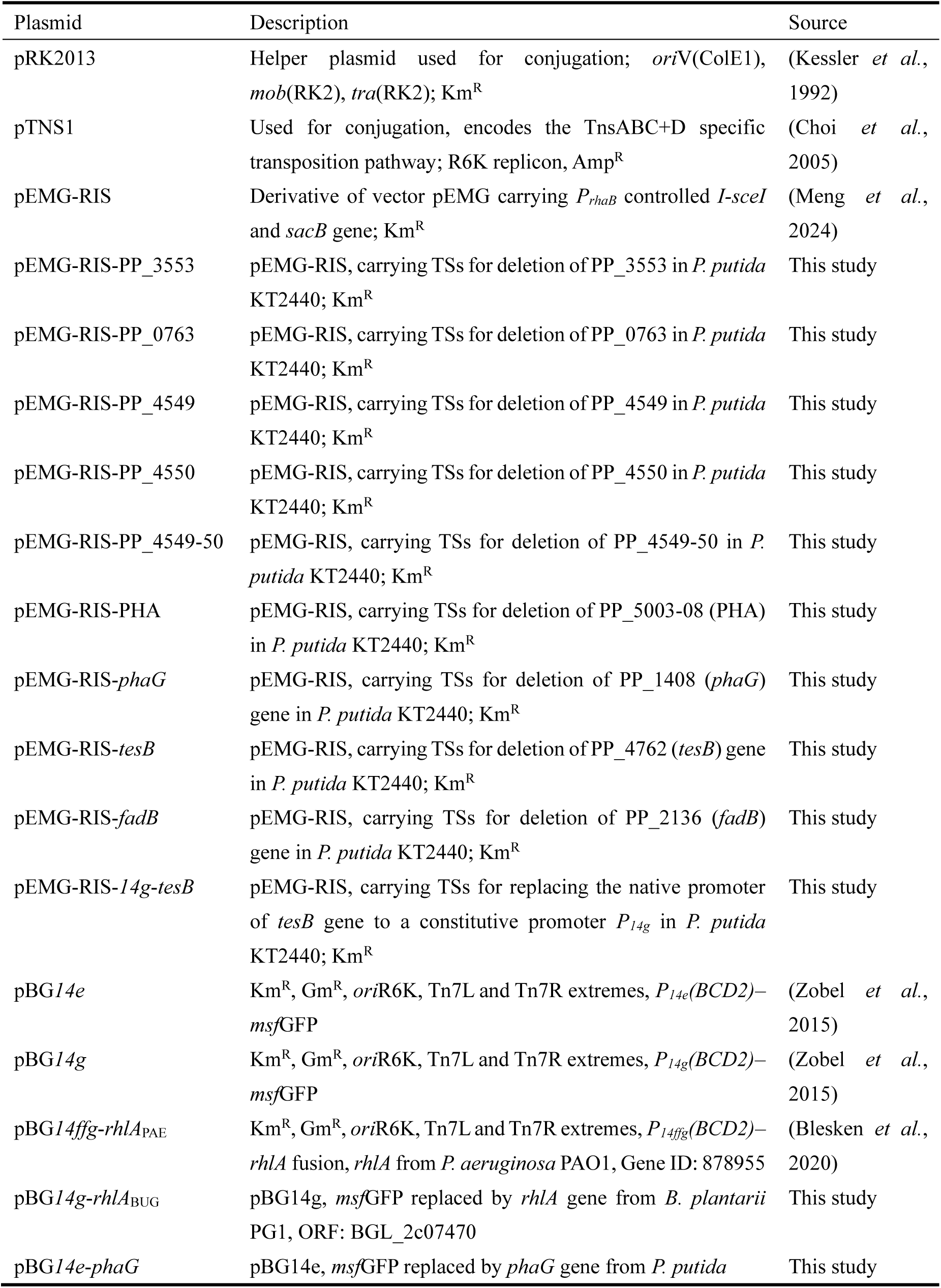
Plasmids used in this study.

Recombinant plasmids in this study were constructed with NEBuilder HiFi DNA Assembly Master Mix (New England Biolabs, Ipswich, Massachusetts, USA). All oligonucleotides were ordered, and all sequencing work was performed at Eurofins Genomics (Ebersberg, Germany). DNA fragments for plasmid construction were amplified using Q5 DNA polymerase, while colony PCRs were performed with One*Taq* 2X Master Mix (New England BioLabs, Ipswich, MA, USA).

### 2.2. Cultivation Conditions

Strains were maintained at -80℃ as cryo-stocks in 25% (v/v) glycerol. For cloning and initial pre-cultures, lysogeny broth (LB) (10 g/L tryptone, 5 g/L yeast extract, 5 g/L NaCl) was used as a liquid medium and as LB agar (1.5% agar). *E. coli* was incubated at 37℃, and *P. putida* at 30℃. After tri-parental mating, *Pseudomonas* transconjugants carrying the desired co-integrate were selected on cetrimide agar. When required, antibiotics were added at the following final concentrations: kanamycin sulfate, 50 mg/L; ampicillin sodium salt, 100 mg/L; and gentamicin sulfate, 25 mg/L.

For 3-HFA production, cultures were grown in a minimal mineral salts medium (MSM) (Hartmans *et al*., 1989), with a modified phosphate buffer adjusted to pH 7.0. In this work, 1× phosphate buffer is defined as 3.88 g/L K_2_HPO_4_ and 1.63 g/L NaH_2_PO_4_ (36 mM phosphate in total). For shake-flask cultivation, MSM with 3× buffer was used together with 2.0 g/L (NH_4_)_2_SO_4_, trace elements, and 10 g/L glucose. The trace elements were added from a 100× stock solution. The final MSM contained (per liter): 10 mg EDTA, 0.1 mg MgCl_2_·6H_2_O, 2 mg ZnSO_4_·7H_2_O, 1 mg CaCl_2_·2H_2_O, 5 mg FeSO_4_·7H_2_O, 0.2 mg Na_2_MoO_4_·2H2O, 0.2 mg CuSO_4_·5H_2_O, 0.4 mg CoCl_2_·6H_2_O, and 1 mg MnCl_2_·2H_2_O. For nitrogen-limited growth, 0.5 g/L (NH_4_)_2_SO_4_ was used to obtain a molar C/N ratio of 44:1 instead of 11:1 at unlimited conditions. For stirred-tank reactor (STR) batch cultivations, a 1× phosphate buffer, 2 g/L (NH_4_)_2_SO_4_, and 10 g/L glucose was applied. During fed-batch fermentation, the feed solution contained 300 g/L glucose, 60 g/L (NH_4_)_2_SO_4_, a 2× phosphate buffer, and a 3.3× concentration of the trace elements.

### 2.3. Transfer Rate Online Measurement

For online monitoring of the oxygen transfer rate (OTR) and carbon dioxide transfer rate (CTR), cultivations were performed in the Transfer Rate Online Measurement (TOM) shaker (Adolf Kühner AG, Birsfelden, Switzerland) using 500 mL Erlenmeyer flasks. Working volumes were varied as indicated; cultures were inoculated to an initial OD_600_ of 0.1. All strains were cultivated at 30℃ and 300 rpm (50 mm orbital throw).

### 2.4. Growth Profiler Cultivation

To monitor the growth of *P. putida* mutants on 3-HFAs, cultures were run in the Growth Profiler (Enzyscreen BV, Heemstede, Netherlands). Strains were cultivated in 96-half-deepwell microtiter plates (CR1496dg; Enzyscreen) with a 0.2 mL working volume at 30℃ and 225 rpm on a 25 mm orbital throw.

### 2.5. Fermentation Setup

Fermentations were carried out in 3.3 L BioFlo 120 stirred-tank bioreactors (Eppendorf AG, Hamburg, Germany) with a 1.2 L working volume, operated using DASware control software version 5.3.1. The system was equipped with online monitoring capabilities, including a dissolved oxygen (DO) probe (VisiFerm DO ECS 225) and a pH probe (EasyFerm Plus PHI K8 225) (both from Hamilton, Bonaduz, Switzerland), as well as Pt100 temperature sensor. The pH was maintained at 7.0 by automatic addition of 4 M H_2_SO_4_ and 4 M KOH. Exhaust gas was passed through a condenser and was analyzed for O_2_ and CO_2_ concentrations using a BlueVary sensor with BlueVis software version 4.65 (BlueSens Gas Sensor GmbH, Herten, Germany). For the batch fermentation, the agitation shaft was equipped with a single six-blade Rushton turbine (53 mm diameter), and DO levels were controlled at 30% by adjusting the agitation speed between 300 and 1,200 rpm with a gas flow of 0.1 to 0.15 vvm. For the DO-based fed-batch fermentation, the agitation shaft was equipped with two six-blade Rushton turbines. It was started with a batch phase, followed by a fed-batch phase after glucose depletion. Feeding was carried out based on the DO signal, between 70% and 30%, with a feed rate of 40 mL/h, which was increased after 2 h to 50 mL/h. With a constant gas flow of 0.1 vvm and an agitation cascade, the DO was controlled until 5 h at 30 % after which a constant agitation rate of 550 rpm was maintained. During the fed-batch phase, the agitation was increased every 1 h by 25 rpm until 950 rpm. Throughout the fed-batch phase, 30 mL of 100× trace element solution was pulsed to the bioreactor.

### 2.6. Determination of Optical Density and Cell Dry Weight

Optical density at 600 nm (OD_600_) was measured with an Ultrospec 10 Cell Density Meter (Amersham Biosciences, Amersham, UK). For cell dry weight (CDW) determination, 2 mL of culture were centrifuged at 15,000 rpm for 5 min at 4℃. The supernatant was retained for substrate analysis. The pellet was washed once with 1.5 mL of ultrapure water, resuspended in 1.0 mL of ultrapure water, and the tube was rinsed with an additional 0.5 mL. The suspension and rinse were then transferred together into pre-dried (65℃, 48 h) and pre-weighed HPLC vials. Vials were dried at 65℃ for 72 h to a constant mass and reweighed to calculate CDW.

### 2.7. Analytics

All analyses were performed on an UltiMate 3000 HPLC system (Thermo Fisher Scientific, Waltham, MA, USA). For quantifying glucose, gluconate, and 2-keto-gluconate, the samples were prepared by centrifuging the cell culture at 13,000 rpm for 5 min and filtering the supernatant with a 0.22 µm syringe filter. Then, an ion-exchange Metab-AAC column (300 × 7.8 mm, 10 µm; ISERA GmbH, Düren, Germany) was used with 5 mM H_2_SO_4_ as the mobile phase at a flow rate of 0.8 mL/min and a temperature of 60℃. Analytes were detected using an internal UV detector at 210 nm and a Shodex RI-101 refractive index detector at 35℃ (Showa Denko, Tokyo, Japan).

For measuring 3-HFAs, a reversed-phase ISASpher 100-5 C18 BDS column (250 × 4.0 mm; ISERA) coupled to a Corona Veo charged aerosol detector (Thermo Fisher Scientific) was used. Mobile phases were acetonitrile (A) and 0.1% formic acid in water (B). The gradient was: 10% A/90% B for 2 min; ramp to 60% A over 4 min and hold for 24 min; increase to 98% A for 2 min; then re-equilibrate at 10% A for 4 min. The column temperature was 40℃. Samples were prepared by mixing culture 1:1 (v/v) with acetonitrile and incubating at 4℃ for at least 4 h before centrifugation and filtration. Commercial 3-HFA standards (C6-C14, even number) were purchased from Larodan (Larodan AB, Solna, Sweden).

Significance analysis was performed by the determination of the standard deviation or standard error of the mean when indicated, followed by an ordinary one-way ANOVA with assumed Gaussian distribution, minimum p <0.05.

## 3. Results

### 1. Blocking 3-HFA catabolism via acyl-CoA ligase deletions

Previous studies have mapped the fatty–acid–activating acyl-CoA ligases in *P. putida* and generated strains that are unable to activate and degrade these molecules (Thompson *et al*., 2020; Valencia *et al*., 2022). We speculated that the same ligases also activate 3-hydroxy fatty acids (3-HFAs) for subsequent catabolism. A combinatorial deletion panel was constructed (PP_3553, PP_0763, PP_4549-50; all single, double, and triple deletion mutants) and growth was assayed on C8, C10, and C12 3-HFAs as sole carbon sources. The mutants displayed chain-length–dependent growth phenotypes, and the triple deletion (PP_0763 and PP_4549-50) abolished growth on all tested 3-HFAs (Fig. 1B–D).

**Fig. 1.**
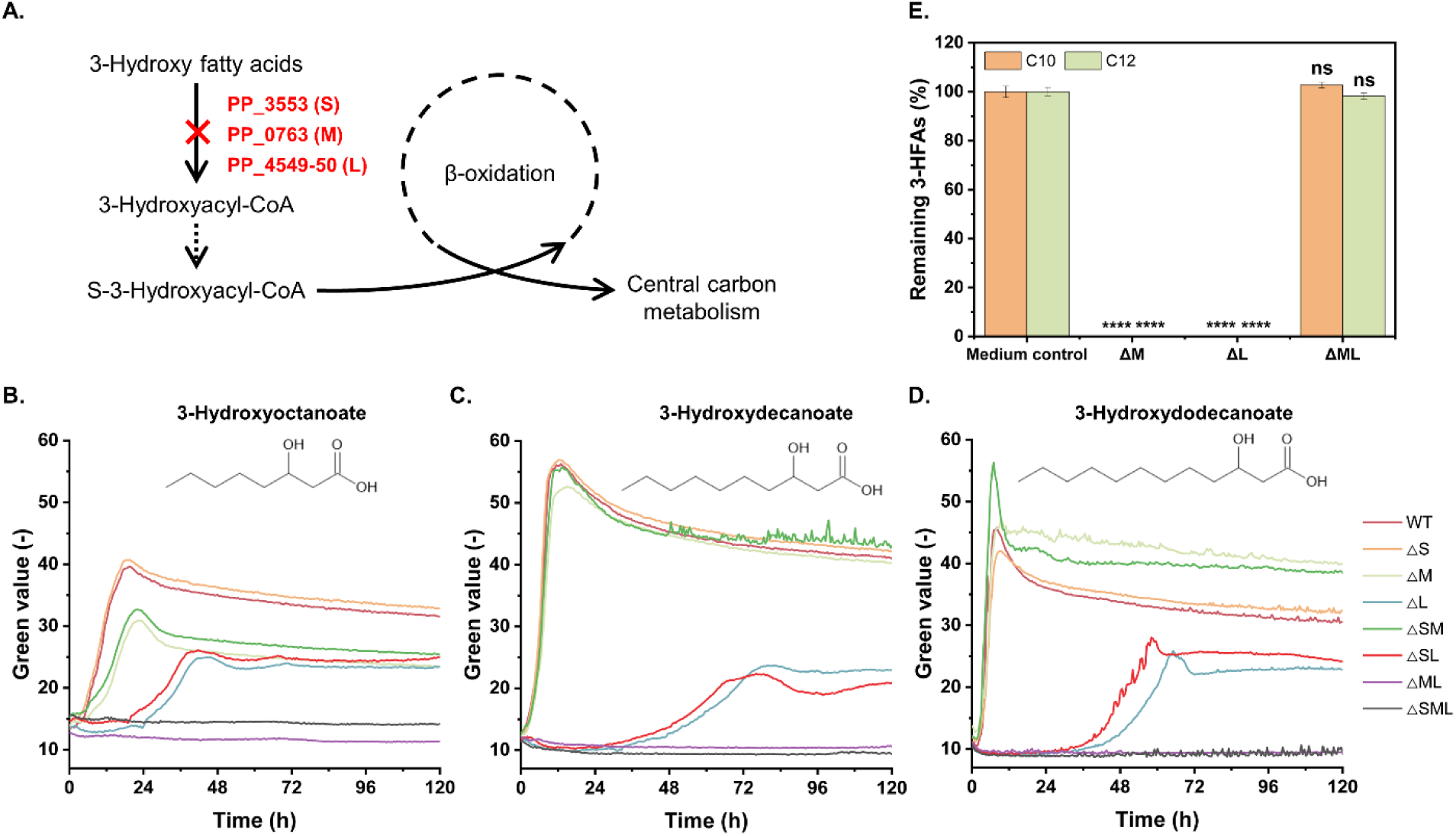
Verification of *P. putida* KT2440 acyl-CoA ligase activity toward 3-HFAs. (A) Combinatorial deletions of acyl-CoA ligases block conversion of 3-HFAs to central metabolism. (B–D) Growth of engineered strains with 1 g/L 3-hydroxyoctanoate (C8), 3-hydroxydecanoate (C10), or 3-hydroxydodecanoate (C12) as sole carbon source. (E) Residual 3-HFAs after growth, quantified and compared with medium control. Error bars indicate the standard error of the mean and **** indicates p < 0.0001 confidence interval of one-way ANOVA analysis. Abbreviations: S, PP_3553; M, PP_0763; L, PP_4549-50. Cultivation conditions: N = 250 rpm; T = 30℃; start OD_600_ = 0.1; n = 2; ns, not significant.

The closer inspection of Fig. 1B–D reveals that for 3-hydroxyoctanoate (C8), deleting either PP_0763 or PP_4549–50 reduced the growth rate, indicating more flexible activation. However, the deletion also results in reduced overall biomass. Deleting PP_4549–50 did not entirely abolish growth but strongly impaired growth on 3-hydroxydecanoate (C10, 3-HDA) and 3-hydroxydodecanoate (C12), whereas deleting PP_0763 had little effect at these lengths. Because PP_4549–50 encodes two ligases (annotated as *fadD*-I and *fadD*-II), single-gene deletions were made to pinpoint their individual function. As shown in Fig. S1, ΔPP_4549 resembled the double deletion ΔPP_4549–50, while ΔPP_4550 behaved like wild type. Thus, for C10 and C12 3-HFAs, PP_4549 is the major ligase, with PP_0763 and PP_4550 providing minor or auxiliary roles. On 3-hydroxyhexanoate (C6), growth occurred only when PP_0763 was intact, pointing to strict control by PP_0763 (Fig. S2). In general, combining ΔPP_0763 (ΔM) and ΔPP_4549–50 (ΔL) was used for subsequent production studies to build on a 3-HFA–non-consuming chassis.

### 2. Investigating native 3-HFA production in *P. putida*

Native 3-HFA formation was examined in the chassis. To avoid interference from mcl-PHA synthesis, the PHA operon (PP_5003–08) was deleted. As shown in Fig. S3, deleting the operon had no apparent effect on 3-HDA formation under nitrogen limitation, and titers continued to rise slowly to ∼72 h, indicating inefficient conversion. Because 3-HDA is the major 3-HFA, it was used as the primary readout for comparing strains and conditions.

Three deletion targets were tested for endogenous 3-HFA formation: deletion of 3-hydroxyacyl-ACP thioesterase *phaG* (often annotated as 3-hydroxyacyl-ACP:CoA transacylase) (Yan *et al*., 2022), deletion of *fadB* (an enoyl-CoA hydratase in β-oxidation), and deletion or overexpression of *tesB* (an acyl-CoA thioesterase reported to release 3-HFAs from R-3-hydroxyacyl-CoA) (Chung *et al*., 2013). Selected combinations were constructed, and 3-HFA production was measured from 10 g/L glucose with either 0.5 g/L (NH_4_)_2_SO_4_ (nitrogen-limited) or 2 g/L (NH_4_)_2_SO_4_. As shown in Fig. 2B, PhaG was the key determinant; its presence or absence dictated whether 3-HFAs accumulated in the ligase-deficient strain. In contrast, *fadB* and *tesB* edits had no apparent effect, implying that hydrolysis of R-3-hydroxyacyl-CoA is not the dominant route to free 3-HFAs. Instead, these data indicate that PhaG acts mainly as a 3-hydroxyacyl-ACP thioesterase rather than a 3-hydroxyacyl-ACP:CoA transacylase under the tested conditions. The consequences of this different activity would be most strikingly for PHA and methyl ketone synthesis, suddenly requiring ligation activity of the monomers (Yan *et al*., 2022).

**Fig. 2.**
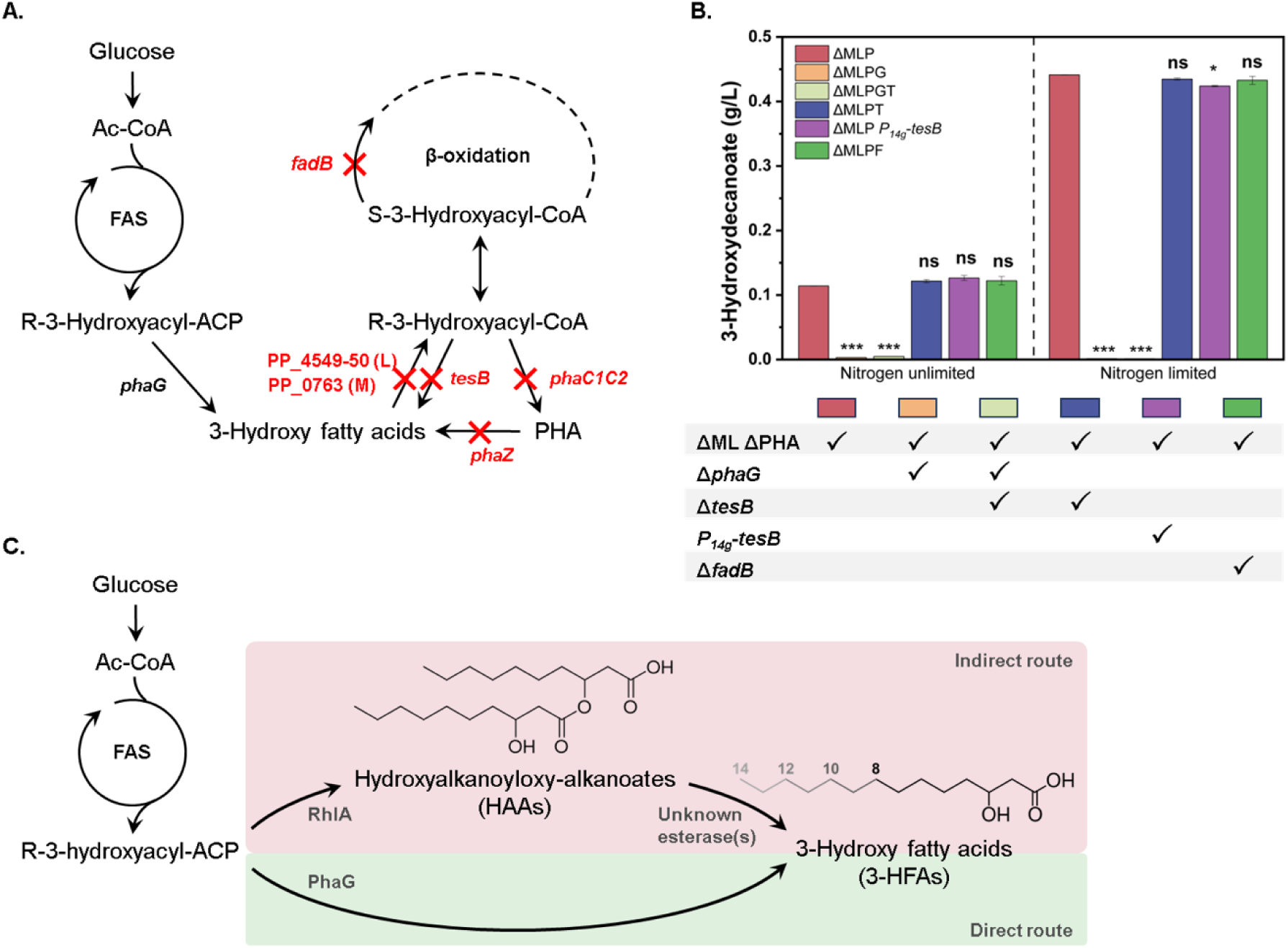
Native and engineered routes to 3-HFAs. (A) Exploiting native release of 3-HFAs from FAS intermediates by PhaG in respective deletion strains. (B) 3-Hydroxydecanoate production by cultivating selected edited strains with or without nitrogen limitation in 100 mL shake flasks in MSM medium supplemented with 10 g/L glucose for 80 h. (C) Two engineered routes: constitutive *phaG* expression and *rhlA*-driven HAA formation with native hydrolysis. Abbreviations: FAS, fatty acid synthesis; HAA, hydroxyalkanoyl-alkanoate; 3-HFAs, 3-hydroxy fatty acids; ΔPHA, deletion of the *pha* operon (PP_5003-08). Error bars indicate the standard error of the mean (n = 2) and * indicates p < 0.05 confidence interval (***, p ≤ 0.001) of one-way ANOVA analysis.

Under nitrogen limitation, glucose was exhausted only after 80 h, and the chassis accumulated 0.44 g/L 3-hydroxydecanoate, indicating that conversion slowed down under strong limitation. To improve efficiency, two alternatives were proposed (Fig. 2C): (i) constitutive *phaG* overexpression to decouple production from nitrogen status; and (ii) *rhlA*-driven HAA formation with native hydrolysis in *P. putida*. Both strategies were evaluated in the following sections.

### 3. Direct 3-HFA production via *phaG* constitutive overexpression

To eliminate native *phaG* effects and ensure constant expression, *phaG* was deleted from its locus and re-integrated at the *attTn7* (between PP_5408 and PP_5409) under the constitutive *P_14e_* promoter (Zobel *et al*., 2015) (Fig. 3A). 3-Hydroxydecanoate production on glucose without nitrogen limitation was then compared across three strains: Δ*phaG*, native *phaG*, and *attTn7*::*P_14e_*-*phaG* (Fig. 3B–D). The native *phaG* strain reached 0.12 g/L, the Δ*phaG* control produced 0.01 g/L, and *attTn7*::*P_14e_*-*phaG* reached 0.55 g/L, a 4.6-fold increase over native and **∼**69-fold over Δ*phaG*, with growth-coupled *P_14e_* promoter activity. The increased expression by *P_14e_* reached titers comparable to previous nitrogen limited conditions, indicating PhaG expression as influential parameter under nitrogen limiting conditions for product synthesis. A minor fraction of 3-hydroxytetradecanoate(s) (C14 and C14:1; 3.2%) appeared in the *phaG* overexpression strain, broadening the chain-length profile compared with *mcl*-PHA production. Testing the strain in LB supplemented with 10 g/L glucose, total 3-HFAs were 1.9-fold higher than on MSM–glucose, while chain-length ratios remained similar, indicating a medium-independent chain length distribution. Because free 3-HFAs and carbon loss by PhaG activity can stress cells, the producer was serially passaged seven times (∼40 generations) in production medium; 95% productivity was retained, demonstrating good genetic stability of *P. putida* ΔMLPG *attTn7*::*P_14e_*-*phaG* (Fig. 3F).

**Fig. 3.**
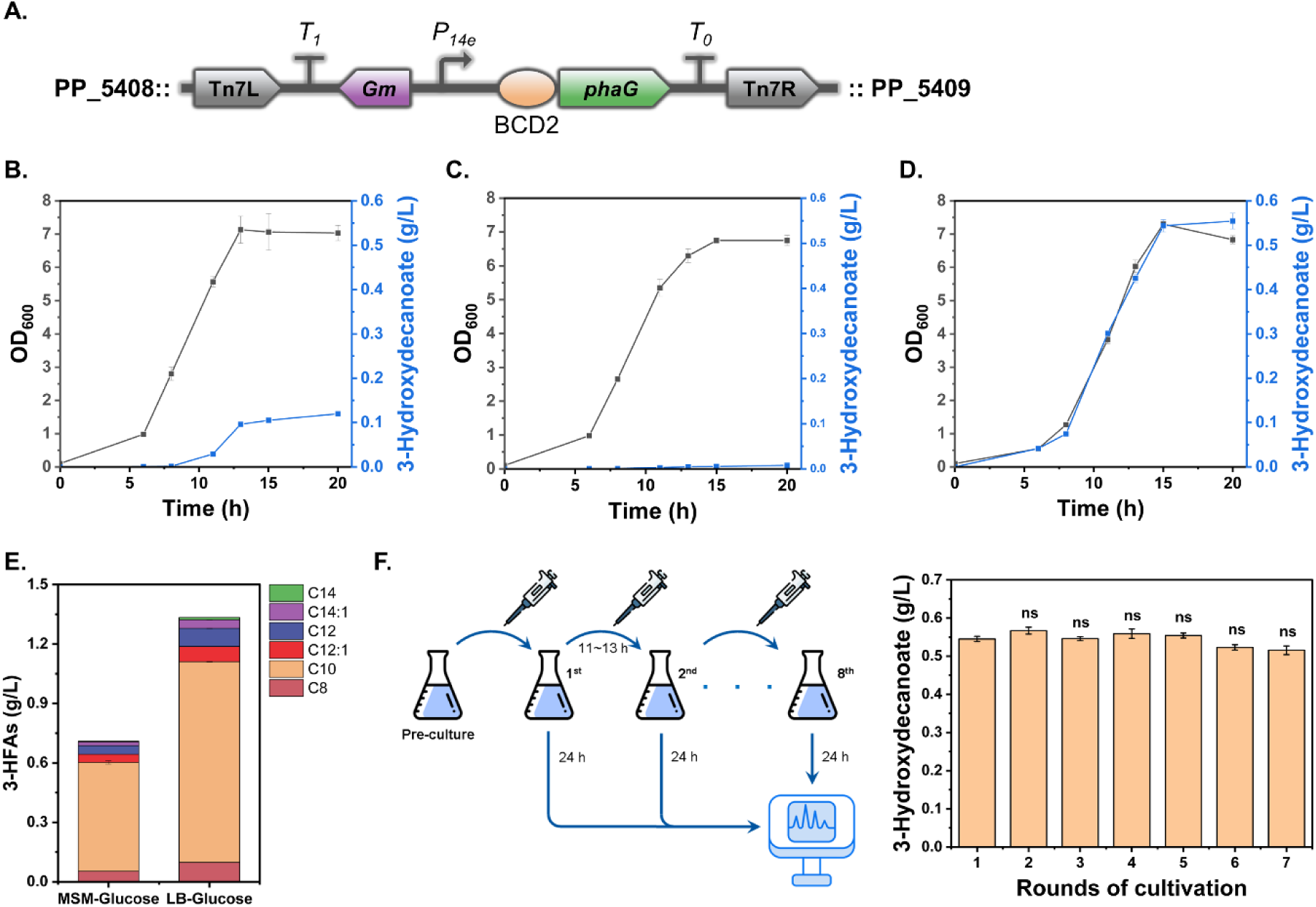
3-HFA production with constitutive *phaG* expression. (A) Chromosomal integration of *P_14e_*-*phaG* at the *attTn7* site in *P. putida* (strain ΔMLPG *phaG*). (B–D) 3-HDA production with native *phaG*, Δ*phaG*, and *attTn7*::*P_14e_*-*phaG*, respectively. (E) Comparison of strain ΔMLPG *phaG* in MSM–glucose vs. LB– glucose (each supplemented with 10 g/L glucose). (F) Genetic stability test of strain ΔMLPG *phaG* for 3-HDA production after seven serial passages in production medium. Error bars represent the standard deviation (n = 3), and ns indicates not significant of one-way ANOVA analysis.

### 4. Incorporating diverse *rhlA* genes for the production of 3-HFAs with distinct carbon chain lengths

To broaden chain-length coverage, heterologous 3-(3-hydroxyalkanoyloxy)alkanoate synthase *rhlA* genes were introduced into the chassis. The *rhlA* from *Pseudomonas aeruginosa* PAO1 (*rhlA*_PAE)_ is known to generate HAA congeners with C8, C10, C12:1, and C12 chains (Tiso *et al*., 2020; Tiso *et al*., 2017; Weiser *et al*., 2022). Prior work investigated *rhlA* variants spanning C8–C18 congener production (Gremer et al., 2020). Accordingly, *rhlA* from *Burkholderia plantarii* PG1 (*rhlA*_BUG_) was selected, and both variants were tested for 3-HFA production via native HAA hydrolysis (Fig. 4A).

**Fig. 4.**
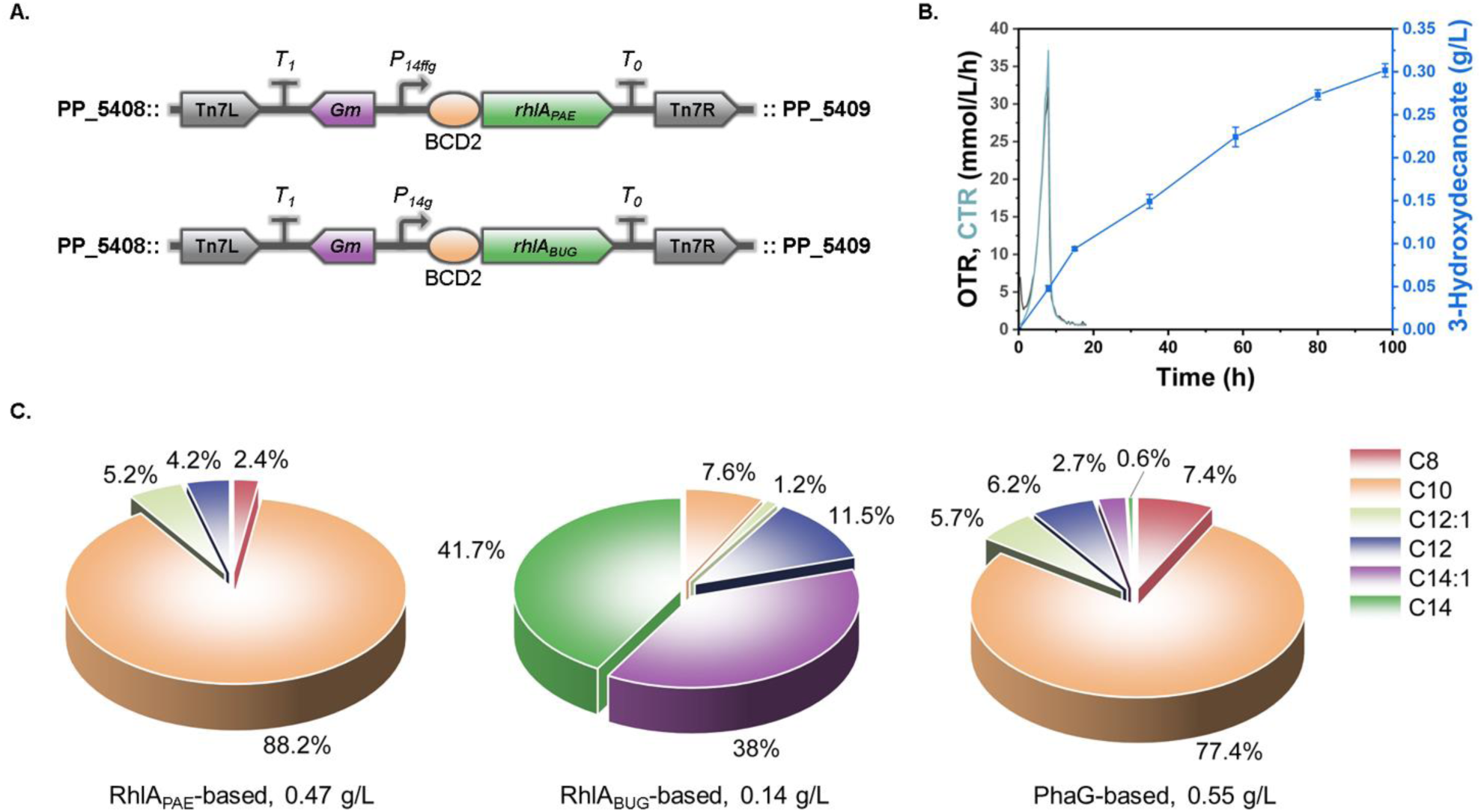
*RhlA*-driven 3-HFAs production via HAA formation and native hydrolysis. (A) Chromosomal integration of *P_14ffg_*–*rhlA* from *P. aeruginosa* PAO1 (*rhlA*_PAE_) and *P_14g_*–*rhlA* from *B. plantarii* PG1 (*rhlA*_BUG_) at *attTn7* in *P. putida*. (B) Time course of 3-HDA release from HAAs by native hydrolysis. The first sample was taken immediately after the OTR dropped, indicating carbon depletion. For this run, the medium only contained 7.5 g/L glucose to avoid oxygen limitation. (C) Composition (%) of 3-HFAs produced by different key enzymes in engineered *P. putida* strains in shake-flask experiments from 10 g/L glucose. left, RhlA_PAE_; middle, RhlA_BUG_; right, PhaG. Error bars are not displayed but remain below 1% (n=3).

Online OTR and CTR monitoring indicated carbon depletion after approximately 10 h (Fig. 4B). In parallel, off-line shake-flask cultures under the same conditions tracked release of 3-HDA from HAAs. Although 3-HDA increased over time, hydrolysis by endogenous esterase(s) was slow, and HAAs were still not fully degraded after 98 h—making this route unattractive for further development at low biomass concentrations and due to a persisting knowledge gap in HAA hydrolysis. Chain-length profiles (Fig. 4C) revealed *rhlA*_PAE_ containing strain produced 0.47 ± 0.02 g/L total 3-HFAs, dominated by C10 (88%) with additional C8, C12:1, and C12; no C14 was detected. The strain with *rhlA*_BUG_ produced 0.14 ± 0.01 g/L total 3-HFAs, enriched in C14 (42%) and C14:1 (38%), with C10 present and only traces of C12/C12:1. These results show that *rhlA* choice controls chain-length distribution and that incorporating diverse *rhlA* genes broadens and specifies the 3-HFA product range.

### 5. Assessing scale-up feasibility in batch and fed-batch fermentations

Following successful shake-flask production via the PhaG pathway, cultivation was scaled to a stirred-tank bioreactor (STR) to assess performance under controlled conditions. First, a batch run was conducted (Fig. 5A–B) with 10 g/L glucose. Airflow started at 0.10 vvm and was increased to 0.15 vvm at 6.6 h to maintain 30% dissolved oxygen. 3-HDA concentration increased simultaneously to the CDW until glucose was fully consumed after 9 h. However, 3-HDA reached only 0.39 ± 0.01 g/L, which is 32% lower than the shake-flask control that was conducted in parallel.

**Fig. 6.**
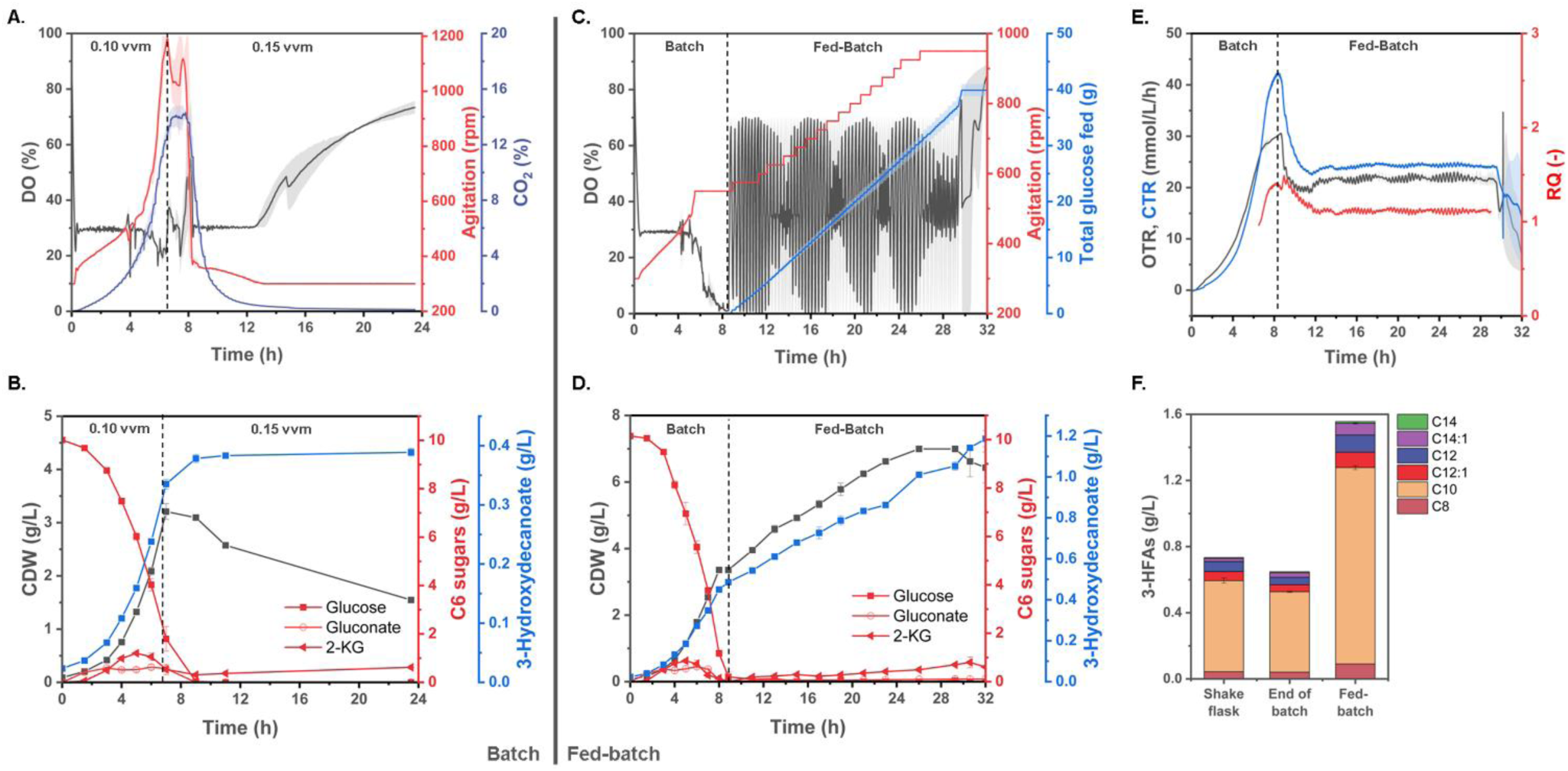
Batch and Fed-batch fermentation of strain *P. putida* ΔMLPG *phaG* for 3-HFAs production. For the batch fermentation: (A) Time courses of off-gas CO_2_, dissolved oxygen (DO), and agitation; (B) Cell dry weight (CDW) and concentrations of glucose, gluconate, 2-keto-gluconate (2-KG), and 3-HDA. For the fed-batch fermentation: (C) Time courses of total glucose fed, dissolved oxygen (DO), and agitation; (D) Cell dry weight (CDW) and concentrations of glucose, gluconate, 2-KG, and 3-HDA; (E) Oxygen transfer rate (OTR), carbon dioxide transfer rate (CTR), and respiratory quotient (RQ). (F) Chain-length composition of produced 3-HFAs in shake flasks, at the end of batch, and at the end of fed-batch in the fed-batch fermentation. Medium: MSM + 10 g/L glucose (shake flasks and fermentations). Fermentation conditions: 3.3 L STR, 1.2 L working volume; 30℃; pH 7.0; air flow 0.10 vvm; starting OD_600_ = 0.2, n = 2. Feeding: DO-based strategy with feed containing 300 g/L glucose, 60 g/L (NH_4_)_2_SO_4_, 2× phosphate buffer, and 3.3× trace-element solution.

Oxygen availability probably explains the titer gap between shake flasks and bioreactors. A TOM shaker experiment with working volume between 4–16% supported this assumption (Fig. S4): 3-HDA titers increased as oxygen availability reduced. At 16% working volume, the titer was 13% higher than at 4% working volume. Accordingly, a second fermentation with a DO-based fed-batch was performed. Oxygen supply was reduced through fixing the agitation rate at 550 rpm from 5 h during the batch phase. This intervention yielded 0.49 ± 0.01 g/L 3-HDA at the end of the batch phase, which is 25% higher than in the first batch fermentation, reducing the shake-flask/bioreactor difference from 32% to 12%. In total, 0.65 ± 0.01 g/L of 3-HFAs were produced with a batch-phase yield of 0.07 g_3-HFAs_/g_glucose_.

After the depletion of glucose, a DO-based fed-batch was initiated at 8.8 h (Fig. 5C, D). Feed pulses were triggered when the DO rose to 70%, and pulsed when the DO decreased below 30%. Until 32 h, 40 g glucose had been supplied per reactor, resulting in 1.56 ± 0.02 g/L total 3-HFAs—equivalent to a fed-batch-phase yield of 0.03 g_3-HFAs_/g_glucose_ and an overall yield of 0.04 g_3-HFAs_/g_glucose_. Throughout the fed-batch phase, the total 3-HFAs increased nearly linearly at ∼0.05 g/L·h, with relatively stable RQ values (Fig. 5E), indicating an almost constant production rate and stable process performance across the run.

## 4. Discussion

We demonstrated an effective, stable, and scalable production of 3-HFAs by constitutive overexpression of *phaG* in a 3-HFA–non-consuming *P. putida* chassis. Under our tested conditions, functional data support PhaG as a 3-hydroxyacyl-ACP thioesterase, rather than an ACP-CoA transferase. The consequences would be wide fetched, if PhaG has only a thioesterase activity from increased energy demand of *de novo* PHA biosynthesis to synthetic coupling of lipid de novo synthesis with beta-oxidation, e.g., for the synthesis of methyl ketones (Yan *et al*., 2022), While in the here presented strains and growth conditions, there is no doubt on the activity of PhaG as a 3-hydroxyacyl-ACP thioesterase, the strains and conditions reported for ACP-CoA transferase activity should be retested.

In defined MSM, total 3-HFAs reached 0.55 g/L in shake flasks and 1.56 g/L in a DO-based fed-batch with a yield of 0.04 g_3-HFAs_/g_glucose_. Additionally, chain-length composition was tunable by introducing diverse *rhlA* genes. These routes and products expand the design space for *de novo* biosynthesis of 3-HFA.

Growth assays on C6–C12 3-HFAs clarified acyl-CoA ligase roles and showed a comparable substrate range with non-hydroxylated fatty acids, as demonstrated by Valencia et al. (2022), who translated the RB-Tn-Seq map into a non-consuming fatty-acid chassis by deleting the medium- and long-chain acyl-CoA ligases (ΔPP_0763, ΔPP_4549-50) in *P. putida* (Valencia *et al*., 2022). 3-HFA accumulation also requires that triple deletion; however, beyond endpoint substrate measurements, residual 3-HFAs were quantified, and growth analysis provides a detailed and visible phenotype readout. The data refine assignments: PP_4549 is dominant for C10–C12 activation; C8 activation is more flexible across ligases; and PP_0763 is essential for C6. Together with prior RB-Tn-Seq mapping (Thompson *et al*., 2020), these results revealed the physiological role of RB-Tn-Seq predicted gene functions from fatty acids to hydroxy fatty acids.

Chain-length composition depends on the chosen gene and route, enabling application-specific tailoring. With the PhaG route, 3-HFAs span C8, C10, C12/C12:1, and C14/C14:1 (Fig. 4C). In the HAA routes, *rhlA*_PAE_ yields C8, majorly C10, and C12/C12:1 (no C14/C14:1). In contrast, *rhlA_BUG_* yields C10, C12/C12:1, and up to 80% C14/C14:1 (no C8) (Fig. 4C). Additional *rhlA* variants—many already profiled across C8–C18 (Germer *et al*., 2020)—can further expand coverage. Besides, structure–function insights from *rhlA* variants that bias monomer length could guide protein engineering of PhaG (e.g., active-site remodeling) to either widen the accessible chain-length range or sharpen specificity toward a desired target product. Moreover, these findings could influence *de novo* mcl-PHA synthesis, as 3-HFAs are the immediate precursors. Broadening the 3-HFA pool should, in turn, translate into more diverse mcl-PHA compositions and properties (Dartiailh *et al*., 2021; Zheng *et al*., 2020).

Fatty acids have antimicrobial activity (Desbois and Smith, 2010), and 3-HFAs do as well (Allen *et al*., 2012; Radivojevic *et al*., 2016), therefore, product toxicity may cap titers. The HAA route can mitigate toxicity because HAAs are less harmful; however, native hydrolysis to 3-HFAs is slow at low biomass. This step could be improved by identifying and overexpressing the native esterase(s) in *P. putida* and performing high-cell-density fermentations, or by applying rapid alkaline hydrolysis (Wang *et al*., 2025). Major drawback is severe foaming when producing via HAAs, since HAAs serve as biosurfactants (Facchinatto *et al*., 2025; Filbig *et al*., 2025; Tiso *et al*., 2021; Wittgens *et al*., 2017).

Scale-up in stirred-tank reactors with DO-based fed-batch confirmed feasibility; however, cultivation and feeding strategies require refinement. Supported by the TOM results and the oxygen-limited batch phase, lower oxygen availability increased titers; thus, DO should be held tightly at a low setpoint. DO-triggered feeding is likely suboptimal because it requires a transient DO rise; a constant or exponential feed that maintains mild oxygen limitation is preferable and merits optimization. A determinant is the periplasmic oxidation of the C-source to gluconate and 2-ketogluconate, which complicates fixed-rate or exponential feeds due to their accumulation. This issue could be minimized by deleting the respective glucose or gluconate dehydrogenases (Chen *et al*., 2024).

To meet the growing demand for bio-based polymers, fuels, and surfactants, this work establishes practical routes for *de novo* production of 3-HFAs. The 3-HFA–non-consuming *P. putida* chassis and modular pathways elucidate metabolism of 3-HFAs and lay a foundation for further engineering. The chassis also enables the re-evaluation of pathways—such as biosurfactant routes—that generate 3-HFAs as transient byproducts but were previously overlooked due to their rapid consumption. By pairing chain-length control with scalable cultivation, the engineered strains enable application-specific monomer supply and advances circular economy manufacturing.

## Author Contributions

Hao Meng: Conceptualization, Data curation, Formal Analysis, Investigation, Methodology, Validation, Visualization, Writing–original draft, Writing–review and editing. Tobias Schwanemann: Conceptualization, Formal Analysis, Methodology, Writing–review and editing. Marie K. Lipa: Methodology, Formal Analysis, Writing–review and editing. Carina V. Michel: Methodology. Jialiang Xia: Investigation. Lars M. Blank: Conceptualization, Funding acquisition, Supervision, Writing–review and editing.

## Supporting Information

The Supporting Information for this article can be found online in the Supporting Information section at the end of this article.

